# Computational anatomy: the cerebellar microzone computation

**DOI:** 10.1101/2024.09.10.612222

**Authors:** Mike Gilbert, Anders Rasmussen

**Affiliations:** School of Psychology, College of Life and Environmental Sciences, University of Birmingham, Birmingham B15 2TT, UK; Department of Experimental Medical Science, Lund University, BMC F10, 22184 Lund, Sweden

**Keywords:** cerebellum, theory, model, computation, anatomy, microzone, network, code

## Abstract

The cerebellum is a large brain structure. Most of the mass and volume of the cerebellum is made up by the cerebellar cortex. The outer layer of the cerebellar cortex is divided functionally into long, thin strips called microzones. We argue that the cerebellar microzone computation is the aggregate of simple unit computations and a passive effect of anatomy, unaided and unlearned, which we recreate *in silico.* This is likely to polarise opinion. In the traditional view, data processing by the cerebellum (stated very briefly) is the effect of learned synaptic changes. However, this has become difficult to reconcile with evidence that rate information is linearly conserved in cerebellar signalling. We present an alternative interpretation of cell morphologies and network architecture in the light of linear communication. Parallel fibre synaptic memory has a supporting role in the network computation.

**Lay summary:** It is not fully understood how the cerebellum represents information and uses it to generate motor outputs, and how outputs are coordinated to control movement. We model the cerebellar network computation, with a focus on the microzone computation, the middle layer of the network. The main proposal is the contentious idea that the core computation is a passive and unlearned effect of neuroanatomy, unit functions are linear, and the computation is mathematically unsophisticated.

To provide functional context, we propose that the topography of input and output of cerebellar motor circuits could be a functional extension of modular cerebellar wiring which automates motor sequences. To illustrate how that could work, we present a simple model of anguilliform (eel-like) swimming.

## 1. Introduction

Most of the mass and volume of the cerebellum is made up of the cerebellar cortex, which is divided into an inner and outer layer, the granular layer and molecular layer. The molecular layer is divided functionally into long, thin strips called microzones [1–3]. Microzones lie at right angles to the main portion of the axons of granule cells, called parallel fibres, which themselves lie parallel to each other and the cerebellar surface. A microzone is populated by a few hundred Purkinje cells. Purkinje cells receive contact in passing from parallel fibres and are the sole outputs of the cerebellar cortex. Firing rates of the output cells of the cerebellum are bidirectionally sensitive to Purkinje cell rates [4]. This provides the main control of output cell rates.

At a mid-range estimate, a microzone is ∼150 µm wide and 15–20 mm long. Purkinje cells are sagittally flattened in the plane orthogonal to parallel fibres but present a large target – an estimated 350,000 parallel fibres pass through the dendritic arbour of a single Purkinje cell [5, 6], and many millions through a microzone. Parallel fibres have a range of ∼3 mm in both directions from the granule cell soma [7, 8], so that a microzone receives parallel fibre input from both sides, from a large region. A microzone accordingly defines a functional network made up of the microzone (the middle layer of the network), the region of the granular layer from which it receives parallel fibre input (the input layer of the network), and the nuclear group it has output to (the output layer of the network).

There is growing evidence that rate information is linearly conserved in signalling between cerebellar neurons [9–12], including synaptic signalling in the molecular layer [13–16], and that cells are adapted for faithful transmission [17–23]. Firing rates linearly code behavioural and task metrics ‘at all levels of the cerebellar circuit’ [24 p.239, citing 30 references]. Neurobiology is highly adapted to control for biophysical and other noise. There are cerebellar examples of this, where cell-to-cell communication is adapted to conserve fidelity of linear transmission of rate information and control for other variables [9–12, 17, 18, 21–23, 25–27].

This paper is an attempt to account for detailed neuroanatomy, network organisation and linear signalling. The main proposals are that i) the core computation is a passive and unlearned effect of neuroanatomy; ii) the network is the physical form of functionally-layered linear unit functions; and iii) the computation is mathematically unsophisticated. Unit functions are attributed to identified anatomical hardware, connected by known anatomy, and evidenced. In this form, we are able to model large cell networks in high computationally-relevant definition. Model parameter values represent coded data and not biophysics. Firing rates are an example, where complex biophysics code a value. We claim that this step allows the model to be biophysically simplified without simplifying the computation.

To provide functional context, we incorporate the network computation in a simple model of anguilliform (eel-like) swimming, a candidate proto-vertebrate wiring plan. We discuss learning in the Discussion section. We propose that parallel fibre synaptic plasticity is in a complementary role to the network computation, with the function of making a pattern-blind, whole-microzone gain adjustment.

While there is a computational element, this is not primarily a computational paper. Many neural network models present a computational mechanism that executes the function of the network. The function is a proposal and becomes an assumption of the model. In practice, these steps may be conflated, but the direction of the argument is function → mechanism (the first two levels of Marr’s three levels of analysis). We work in reverse. We infer a mechanism from the evidence and then infer function from the mechanism.

This is significant. In our approach, the simulation provides a test of the performance of the system. The purpose is to demonstrate proof of concept, as opposed to being the form of a mathematical generalisation. ‘Proof’ does not refer to mathematical proof, but a practical demonstration, using examples.

We do not propose or claim to be mathematically interesting. The sophistication of the system lies in its biological implementation, isolating the relationship of parameters that code information and controlling for other variables, including biophysical noise. A note on terminology: a layer of a network has a different meaning to layers of the cerebellar cortex. For example, a microzone, the middle layer of a network, is a division of the molecular layer, a stratum of the cerebellar cortex.

Sections 2 and 3 describe the derivation of parallel fibre input to a microzone. Section 4 describes the proposed microzone computation. Section 5 describes a simple system which incorporates the network computation.

## 2. A network of linear functions

### 2.1 Computational anatomy

We have previously simulated the granular layer mechanism which converts mossy fibre signals into granule cell and therefore parallel fibre signals [28]. In the granular layer model, the granular layer is divided functionally into long thin strips – ‘ranks’ – which mirror microzonal organisation. We envisage that anatomy is the physical form of a network of computational units. A unit takes inputs and returns output. Inputs and output represent values (that is, numbers). Physically, a unit is the site of biophysical events which relate inputs and output – the unit function. Unit functions are linear – output is a linear function of inputs. A review of evidence of linear signalling in the granular layer appears in Supplementary materials.

The physiological form of a unit depends on the layer/unit type. Units are physically intermingled in both a rank and in a microzone, but organised functionally in layers. The outputs of a layer provide the data sampled (i.e., received as inputs) by units in the next layer. Computations run continuously, so unit outputs are constantly refreshed – i.e., analogue.

All layers of unit functions span the network from side to side (because the population of a layer occupies either the volume of a rank or a microzone). Information is coded collectively in the concurrent outputs of the whole population of a layer. It is coded as a frequency distribution, so that a whole layer codes information that is both indivisible (cannot be subdivided into smaller units of information) and yet coded in any random sample of unit output values.

### 2.2 How are units connected?

Units are connected by anatomy, so we know what the connections are, because anatomy has been reported in detail. The number of inputs to a unit is usually anatomically randomly variable in a reported range. In some cases, we simulate that by generating the number with a distributed probability, and in others we use the mean.

It is unnecessary to know which individual cells contact each other because (biologically and in the model) contact is at random inside topographically defined dimensions. This has the asset for modelling that a target population can be represented by randomly sampling data received as input to a location.

Biologically, a location is a volume with spatial dimensions. *In silico*, we can represent that as a population of unit functions. We know from anatomy (or can derive) the size of unit populations, population ratios, and convergence and divergence ratios. These provide model parameters and the values they take.

### 2.3 How does that ‘look’ in the granular layer?

Stated in short form, the input layer of a network receives input from what may be around 80,000 mossy fibres. Inputs are passed through a chain of iterative random samplings (simplified: Golgi cells sample mossy fibre rates and glomeruli sample Golgi cell rates) to calculate inhibition of granule cells at each of 700 glomeruli per field, in 4,100 fields (into which we nominally divide the input layer of the network; Fig 1). Outputs are passed to the granule cell computation to derive the independent outcome of comparing inhibition and randomly sampled excitation of each dendrite of around 9,000 granule cells per field, field by field, to obtain the number that fire in each field, and the rates they fire at on a normalised scale. The proportion of granule cells which fire is well-predicted by probability, which is controlled by the relationship of inhibition and the mean of mossy fibre rates. At collective level, granule cell firing rates are well-predicted by mossy fibre rates, and have a focused range centred on the mean.

**Fig 1.**
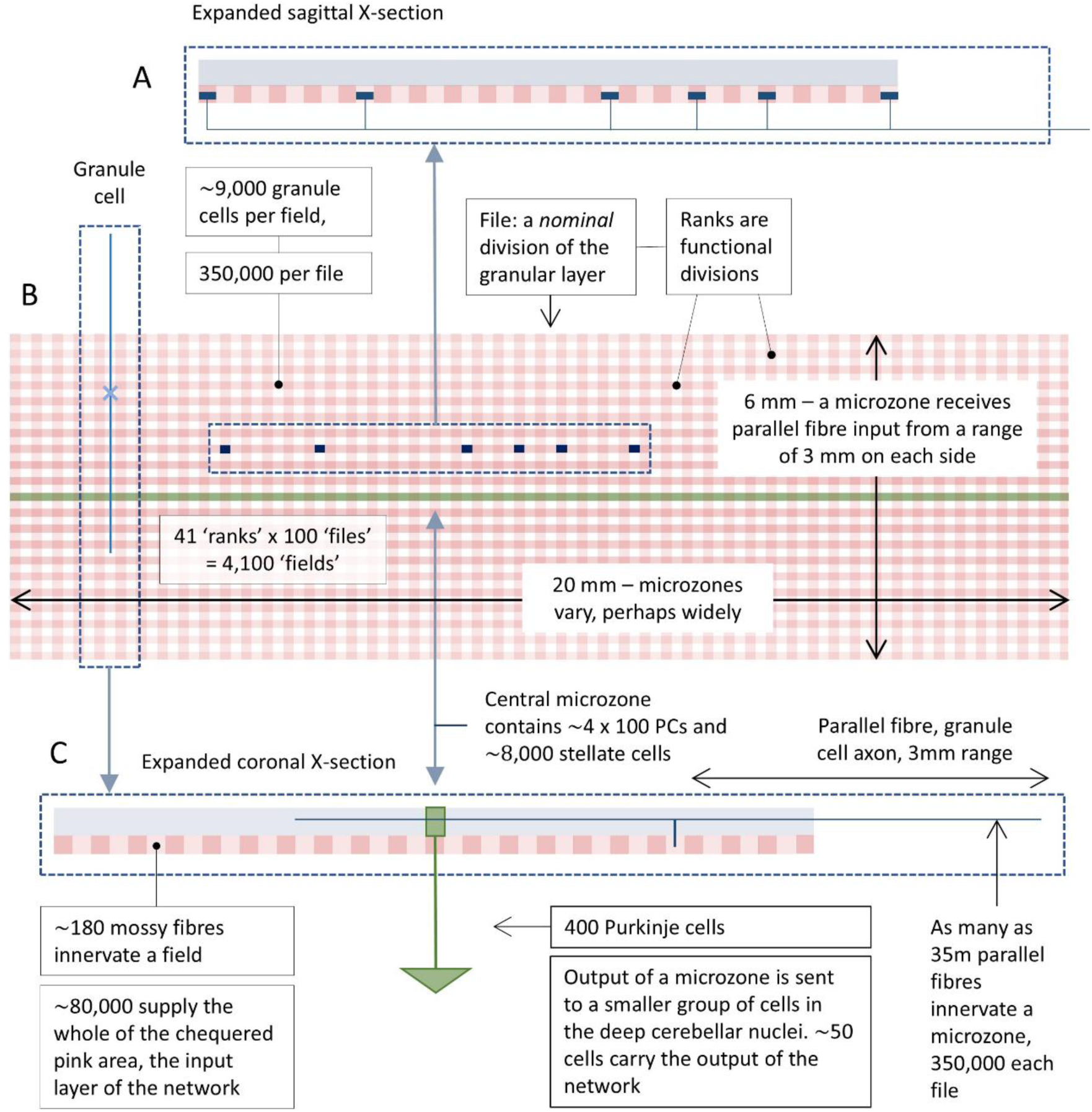
Schematic of a network. Schematic of a region of the flattened cerebellar cortex made up of the input (pink) and middle (green) layers of the cerebellar network, showing relative dimensions of functional and nominal subdivisions. **A** Light blue: molecular layer; pink: granular layer; chequered pattern: files in cross-section; dark blue blocks: the dimensions of the volume occupied by an average mossy fibre terminal cluster. In reality, clusters do not fit neatly into fields or files, and terminals are randomly intermingled with terminals of other mossy fibres. There are around 700 terminals in total per field; a single mossy fibre contributes less than 1%. **B B**oth the molecular and granular layers are functionally divided into long, thin sagittal strips defined by the way input maps to the cerebellum. Granular layer strips are ‘ranks’. A microzone receives parallel fibre input from around 40 ranks, 20 each side. ‘Files’ are nominal divisions, simply a modelling device to quantify the granular layer computation and in doing so to derive the number of active granule cells that provide input to a microzone sector, and the rates they fire at. Only a single microzone is shown. Each microzone defines its own network, so the input layers of near neighbouring networks overlap very greatly. **C** The output of a microzone is the continuous firing of Purkinje cells which converge onto an associated nuclear group.

### 2.4 In this paper, the form of the granule cell/parallel fibre code is an assumption

We use the granular layer mechanism to generate parallel fibre input to the microzone computation. Previously, we modelled a single rank [28]. In this paper, we extend that to the larger region (around 40 ranks) that supplies parallel fibre input to the microzone that equally bisects the region viewed from the cerebellar surface.

In this paper, the workings of the granular layer mechanism are effectively an assumption. Recoding in the granular layer generates a constantly changing pattern of active parallel fibres but with invariant features, regardless of the information it contains. Information is coded at collective level, in the joint activity of cells which are active at the same time. More exactly, it is coded in the frequency distribution of firing rates where active cells intersect a sagittal plane. Firing rates are randomly distributed among active parallel fibres and active parallel fibres are a random subset of the general population. Accordingly, any random sample of firing rates is (at any moment) a random sample of the same distribution. Functionally, a microzone is the physiological form of a sagittal plane. So, because all sectors of a microzone receive a large number of active inputs at any time, all sectors receive the same code at the same time.

### 2.5 The key importance of mossy fibre morphology

#### 2.5.1 MOSSY FIBRE TERMINAL MORPHOLOGY, ‘RANKS’ AND ‘FIELDS’

Mossy fibre morphology has a computational effect, we submit. Mossy fibres give rise to sagittally-extending collaterals, the direction of the long axis of microzones. Collaterals branch terminally, and each branch ends in a cluster of terminals, so that a single cell terminates in a spaced-out row of terminal clusters which are always lined up in the same direction (parallel to microzones but in the granular layer) [29–32]. Clusters are separated by a randomly variable distance typically not less than 200 µm [29, 30] measured from edge to edge of clusters (so they are separated by a greater distance measured from centre to centre). Terminals are small compared to the average volume occupied by a cluster, and intimately and randomly intermingled with terminals of other mossy fibres.

Accordingly, mossy fibres do not terminate at a point but a region with dimensions: the field occupied by a terminal cluster. A single mossy fibre gives rise to several terminal branches and each of those to several terminals, such that a sagittal strip of the granular layer receives multiple copies of every signal at each of multiple locations. The dimensions which enclose a cluster are variable (partly because it depends on the number of terminals in a cluster), averaging 200 µm sagittally x 150 µm mediolaterally [30]. A sagittal row of 100 cluster fields (‘fields’) has the dimensions of a rank viewed from the cerebellar surface (Fig 1). Fields are nominal, a modelling device. Ranks are functional divisions – the minimum dimensions at which the cerebellum makes any effort to tell signals apart, we submit.

#### 2.5.2 TERMINAL BRANCHING

Mossy fibre terminal morphology has important computational significance (for a full discussion see [28]). Terminal branching of mossy fibre collaterals simulates independent random sampling, by fields, of input rates to a rank, also called sampling with ‘replacement’. If each mossy fibre could terminate in only one field, a mossy fibre ‘sampled’ by one field clearly cannot also be sampled by any others. So, samples are not independent. Termination by a single mossy fibre in a random number of terminal branches, at multiple locations, removes that restriction.

In theory, sampling with replacement would mean any number of fields can receive the same signal, up to all of them. Clearly this is not true – in reality there is an anatomical limit. However, even if there was real replacement, the number of repeat samplings is constrained by a probability distribution. If the number of terminal branches per mossy fibre has the same probability distribution, branching has the same result as if firing rates received by each field in a rank were in fact a random sample with replacement of firing rates received by the rank as a whole.

#### 2.5.3 TERMINAL CLUSTERING

Mossy fibre terminal clustering simulates independent random sampling, by individual cells, of input rates to a field. The principle is the same as with terminal branching, but at local scale. Mossy fibre terminal clustering, and random intermingling with terminals of other mossy fibres, means that contact on any particular cell does not mean other cells cannot also receive the same signal. Again, to approximate independence, it is only necessary for the number of repeat selections of a signal to have the same probability distribution as if there was actual replacement.

#### 2.5.4 THE COMBINED EFFECT OF TERMINAL BRANCHING AND CLUSTERING

The combined effect is a biological facsimile of simultaneous random sampling with replacement by *individual* granule cells and Golgi cells of firing rates received as input to *a rank*, notwithstanding that single cells individually receive fixed contact from a tiny fraction of mossy fibres. Note that it is the population of mossy fibre *rates* that are hypothetically sampled with replacement, and not *signals.* The population of signals includes copies of rates because of branching and clustering.

As a result, all granule cells in a rank randomly sample, at any time, the same distribution, and information represented by the output of a rank – i.e., granule cell rates – is coded in any random sample of signals [28]. Since all sectors of a microzone receive parallel fibre input from all the same ranks, microzone-grouped cells all simultaneously randomly sample, with simulated replacement, the whole population of parallel fibre rates received by that microzone. Independent random sampling by topographically-defined cell groups is the cornerstone of the network computation.

#### 2.5.5 WHY IS THAT KEY?

Say a population of values (with any frequency distribution) is randomly sampled with replacement. There are *n* samples, sample size *m*. If we take the mean (or sum) of each sample, and *n* is large, the sample means have a normal or near-normal distribution, by the central limit theorem. (This works even if *m* is a range.) The new distribution is centred on the same mean as the sampled distribution, but narrower.

Large values of *n* and *m* give a nearly-normally-shaped new distribution. Modest values give a normal-tending distribution. However, if the new distribution is then itself independently randomly sampled, and the process is repeated, the results become more focused, and the relationship of the means of successive distributions is well preserved. We will argue that the cerebellar microzone provides (part of) the physical hardware for the cerebellar network to distil output from input in this way, noting that at each step the relationship of the means is adjusted by a constant.

A precis of the ideas in this section and a schematic discussion of merits and limitations appears in Figure 2.

**FIG 2.**
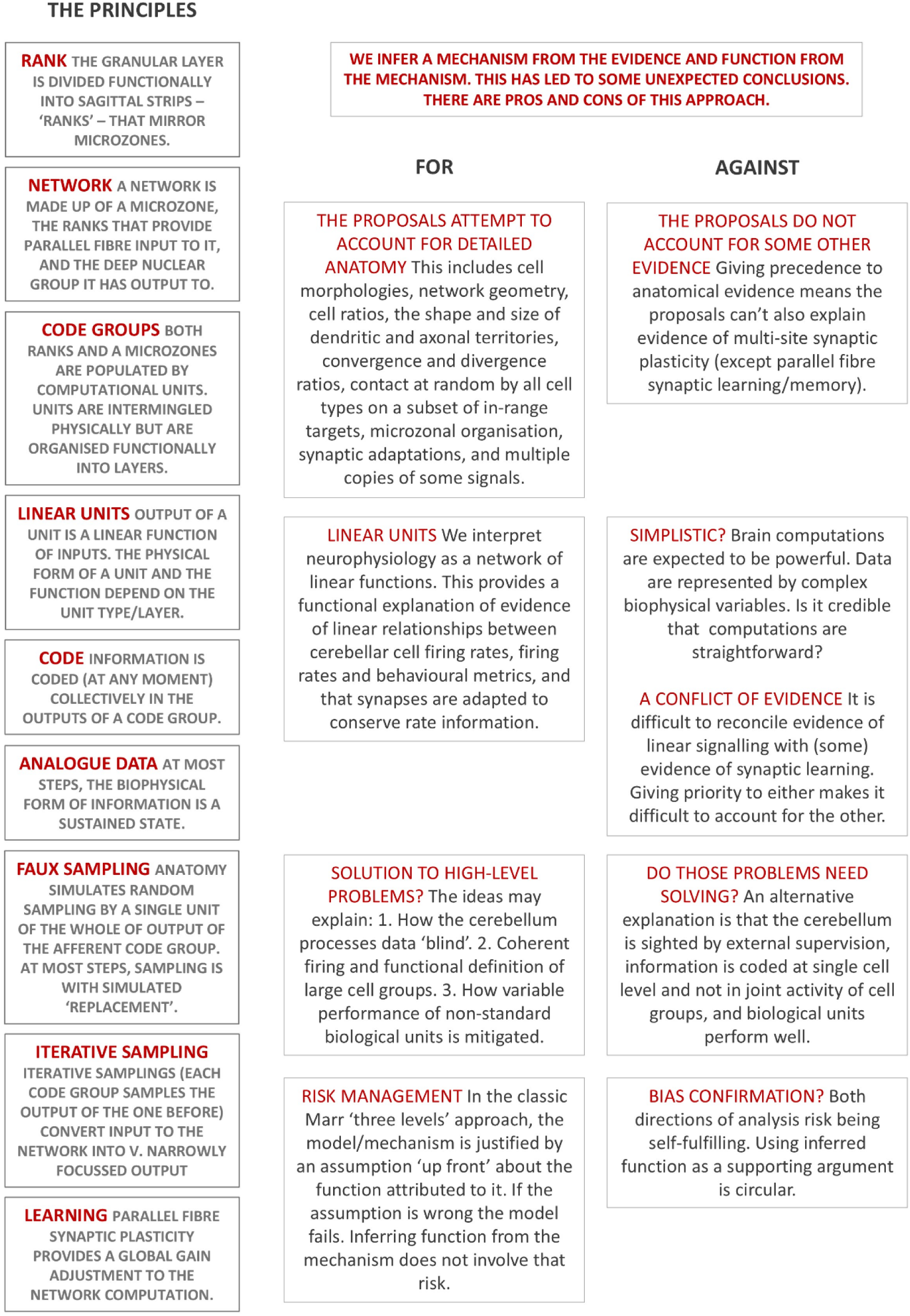

### 2.6 Derivation of mossy fibre rates used in the simulation

We use the simulation to test if an anatomical computation is feasible. For that, it is necessary to generate input to the system that reflects mossy fibre terminal branching and terminal clustering, and also to generate input to each field which reflects the fact that cluster size is randomly variable and most clusters straddle field limits, so that some terminals fall inside and some outside. The derivation of mossy fibre rates received as input to a rank and to fields is described in Supplementary materials.

## 3. Regulation of parallel fibre activity

### 3.1 A case for sparse code (small fraction)

To simulate the computational effect of random sampling of parallel fibre signals, we need to know or derive sample sizes (the number of inputs to each type of unit), and therefore in turn what fraction of parallel fibres is active at any time. There is a long-standing idea that the fraction of parallel fibres that are active at any time is regulated [33], fixed [34], and low (termed sparse code). We disagree with the theoretical reasons, but agree the code is sparse, on these grounds.

1) Low density is stable (by our calculation – Fig 3). Randomly fluctuating density would be noise. Stability may be enhanced by feedback via granule cell ascending axon contact on Golgi cell basal dendrites [19] and parallel fibre contact on Golgi cell apical dendrites, acting partly through gap junctions [35].
2) Low density is energy efficient. Granule cells are very numerous, and they are not myelinated (so spike propagation is costly).
3) The reported number of coincidentally active inputs to a stellate cell is in single figures [36], consistent with low density of co-active parallel fibres (see Table 1).
4) It is thought that the pattern of active and inactive parallel fibres, and the pattern in which firing rates are distributed among active cells, are decorrelated – effectively, quasi-randomised. Targets receive contact from a subset of passing parallel fibres – Purkinje cells receive contact from 1 in 2 [37] and stellate cells from something like 1 in 80 (Section 4). Low density enhances practical decorrelation: if decorrelation is not perfectly random – as it may not be [38] – low density increases the randomness of the pattern of synaptic activations received by parallel fibre targets.

**Fig 3.**
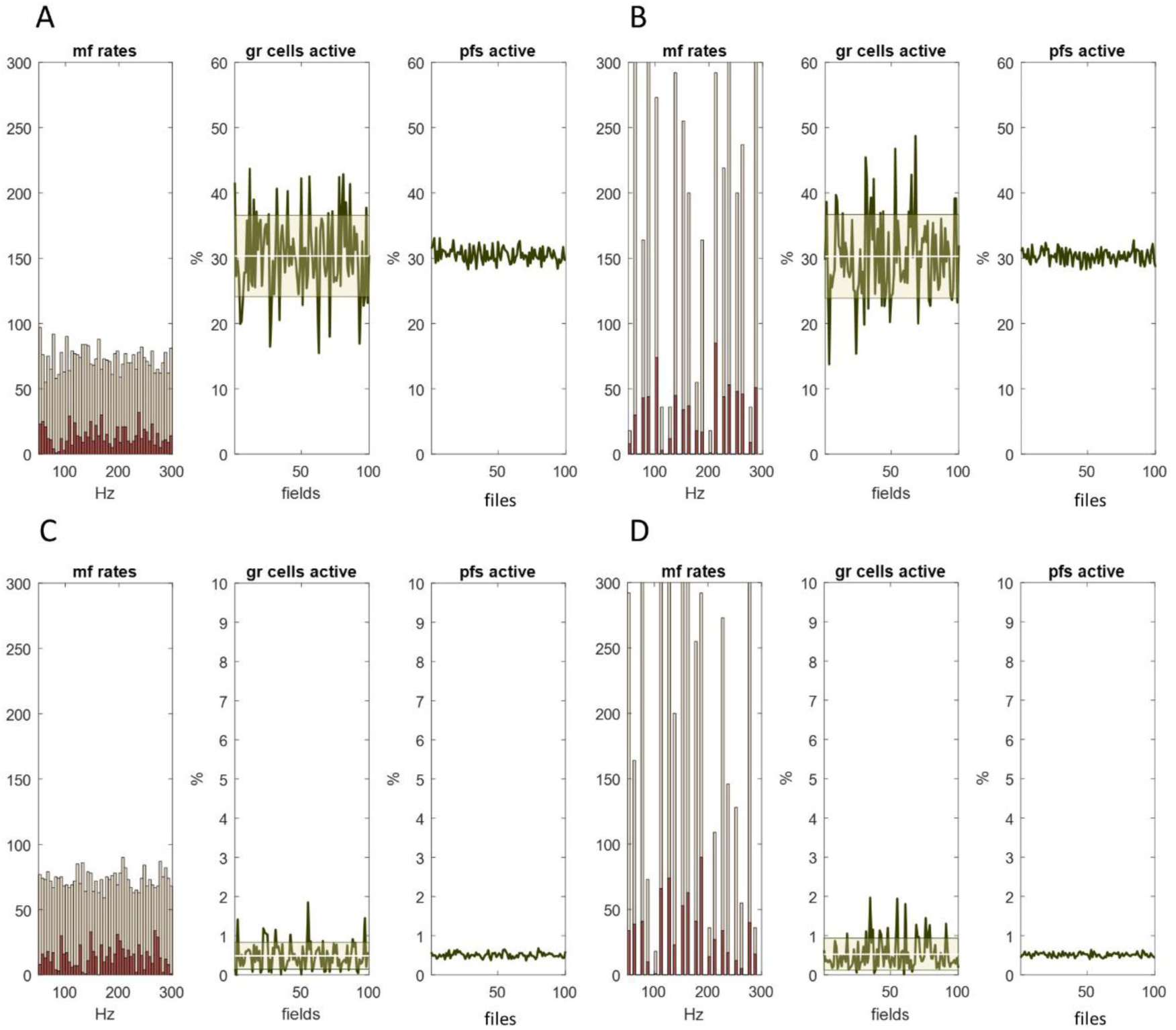
Recoding in the granular layer regulates the fraction of active parallel fibres. Columns 1 and 4. Firing rates received as input to the granular layer region that supplies parallel fibre input to a microzone. Data represent inputs to a single rank, selected at random. Inputs to each rank were independently randomly generated with a uniform and discontinuous distribution, respectively. The method of generating mossy fibre rates received by each field (in all ranks) is described in Supplementary materials. Data represent a short period, effectively a moment of time. **Columns 2 and 5.** The percentage of granule cells that fire in each field of a single rank, again selected at random. **Columns 3 and 6.** The percentage of granule cell that fire in each file, therefore also the percentage of active parallel fibres. **Top row.** ∼30% of granule cells are active. **Bottom row**. Golgi cell inhibition of granule cells is adjusted by a constant so that a lower fraction of granule cells fire, around 0.5%. y axes are scaled to the data (so the contraction of the data range in the bottom row is even greater than it appears). The shape of the sampled distribution (columns 1 and 4) does not affect the results, suggesting that a changing permutation of input to the system at time-varying rates does not alter the fixed fraction of active parallel fibres.

**Table 1.** The probability, *p*, that a stellate cell receives contact from *x* active parallel fibres. Table 1 shows the distributed probabilities of the number of active inputs to an inner-level stellate cell.

| $x$ | 0 | 1 | 2 | 3 | 4 | 5 | 6 | 7 | 8 | 9 | 10 | 11 | 12 |
| --- | --- | --- | --- | --- | --- | --- | --- | --- | --- | --- | --- | --- | --- |
| $p$ | .0066 | .0331 | .0836 | .1403 | .1762 | .1765 | .147 | .1047 | .0651 | .0359 | .0178 | .008 | .0033 |

For these reasons, we calibrated the granular layer model to convert input to the system into a low fraction of active granule cells. We chose 0.5% active because it has support from (as far as we know) the only estimate of the number of co-active inputs to a stellate cell (per Section 4). We suspect it may be higher – perhaps 1% – but a very low fraction is a more rigorous test of our proposals (because sample sizes are smaller).

### 3.2 Simulated regulation of the fraction of active parallel fibres

As noted, we extended the granular layer model to simulate recoding of input to a network. One of the outputs of the model is the number of active granule cells per unit volume of the granular layer. We use fields as the unit of volume. A mediolateral row of fields is a ‘file’ (Fig 1). The input layer of a network is accordingly crossed in one direction by 41 ranks (20 either side of the middle microzone and one directly beneath) and at right angles by 100 files, making 4,100 fields.

The number of active granule cells in each field gives the number in each file. The number in a file is equal to the number of active parallel fibres in that file that converge on the middle microzone. The results are shown in Figure 3. Local regulation of the number of active granule cells in each field in a rank (Fig 3 columns 2 and 5) becomes still tighter regulation of the number of active parallel fibres in a file (Fig 3 columns 3 and 6), a statistical effect of cerebellar cortical architecture. Density is independent of the frequency distribution of mossy fibre rates (Fig 3 columns 1 and 4). At low density of active granule cells, the regulated density of active parallel fibres is very strict.

Physiological regulation may be still stricter, that is, the number of active granule cells in each field may not vary as much as it does in the simulation. Local feedback via the granule cell ascending axon [19, 39] may condense the physiological range of field-to-field variation. Feedback via parallel fibres [34, 35] may in addition reduce the range of file-to-file variation. These are not represented in the simulation.

So, by our calculation, data are not corrupted by noisy variation of the fraction of active parallel fibres, because the fraction is strictly regulated. The primary regulator is the granular layer computation. This has functional significance in two ways. The first is that it means the distributed probability is unchanging of the number of active inputs received concurrently by parallel fibre targets. The second is that it means Purkinje cells receive a fixed number (but high turnover) of active inputs, so that learning can influence Purkinje cell firing by controlling the number of signals that Purkinje cells receive (vs learned changes of graded synaptic weights). We develop this idea in the Discussion.

## 4. The microzone computation

### 4.1 Stellate cells

#### 4.1.1 SAMPLE SIZE

Purkinje cells receive inhibition from interneurons, stellate cells (some vertebrate groups have a second phenotype not included here). The stellate cell territory is sagittally flattened in the plane orthogonal to parallel fibres. Stellate cells form planar networks [40] that occupy the space (about 40 µm) between Purkinje cells [41]. The stellate cell dendritic territory varies with the depth of the cell body in the molecular layer. For superficially located (‘outer-level’) cells it is around 80 x 80 µm. At deeper level (‘inner level’) it is larger, around 120 x 120 µm [41 pp.217-221]. The number of parallel fibres which pass through an inner-level stellate cell territory is given by *a*/((*b* x *c*)/120^2^), assuming parallel fibres are uniformly distributed, where *a* is the number that pass through a Purkinje cell territory (350,000 [6]), and *b* and *c* are the dimensions of the Purkinje cell territory in the same plane (200 x 300 µm), giving ∼84,000. If 0.5% are active in the general population, around 420 of those are active.

It has been estimated that a stellate cell receives contact from several hundred parallel fibres [41, 42], which we take as 1,000. Working with these estimates, 1 in 84 make contact (i.e., 1,000 out of 84,000), so the probability that an active cell makes contact is 1/84 = ∼0.0119. Accordingly, contact by *x* (out of 420) active cells has a probability given by

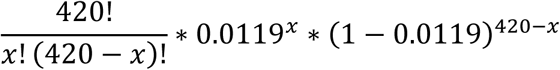

Probabilities are shown in Table 1 for the range *x* = 0 to 12, where *x* is the number of active parallel fibres that make contact and *p* is the probability of that number. The distribution of probabilities is in good agreement with the modest number – single figures – suggested by ‘two to eight substantial EPSPs’ [36 p.9628].

#### 4.1.2 SIMULATED SAMPLING AND INTEGRATION

We notionally divided the middle layer of a network (the middle microzone in Fig 1) into ‘sectors’. Sectors are the same size as fields viewed from the cerebellar surface so that a sector and the field below it together form a column of the cerebellar cortex. Purkinje cells and stellate cells in a sector randomly sample parallel fibre rates in the file that intersects a microzone at that location. (Like fields, sectors are notional and do not represent functional segregation of data.)

The granular layer model returns the firing rates of granule cells in each field, and therefore the frequency distribution of parallel fibre rates in each file, and accordingly rates received by each sector. There are an estimated 16 stellate cells for each Purkinje cell [43], meaning 80 stellate cells per sector (there are four Purkinje cells per sector so five layers of stellate cells are needed to inhibit them from both sides). Sample size was randomly generated with the probabilities in Table 1. This gave the number of active inputs, and input rates, to each stellate cell in each sector.

Molecular layer interneurons, which include stellate cells, reflect ‘granule cell input with linear changes in firing rate’ [15 p.6]. We assume that stellate cell firing is a linear function of the mean of input rates, which we take as the mean itself (the sum works as well). As the physiological function is not known, and our purpose is only to demonstrate a statistical relationship of data layers (i.e., the outputs of units functions functionally organised into layers), and not to calculate actual firing rates, we use a normalised scale. The results are shown in Figure 4.

**Fig 4.**
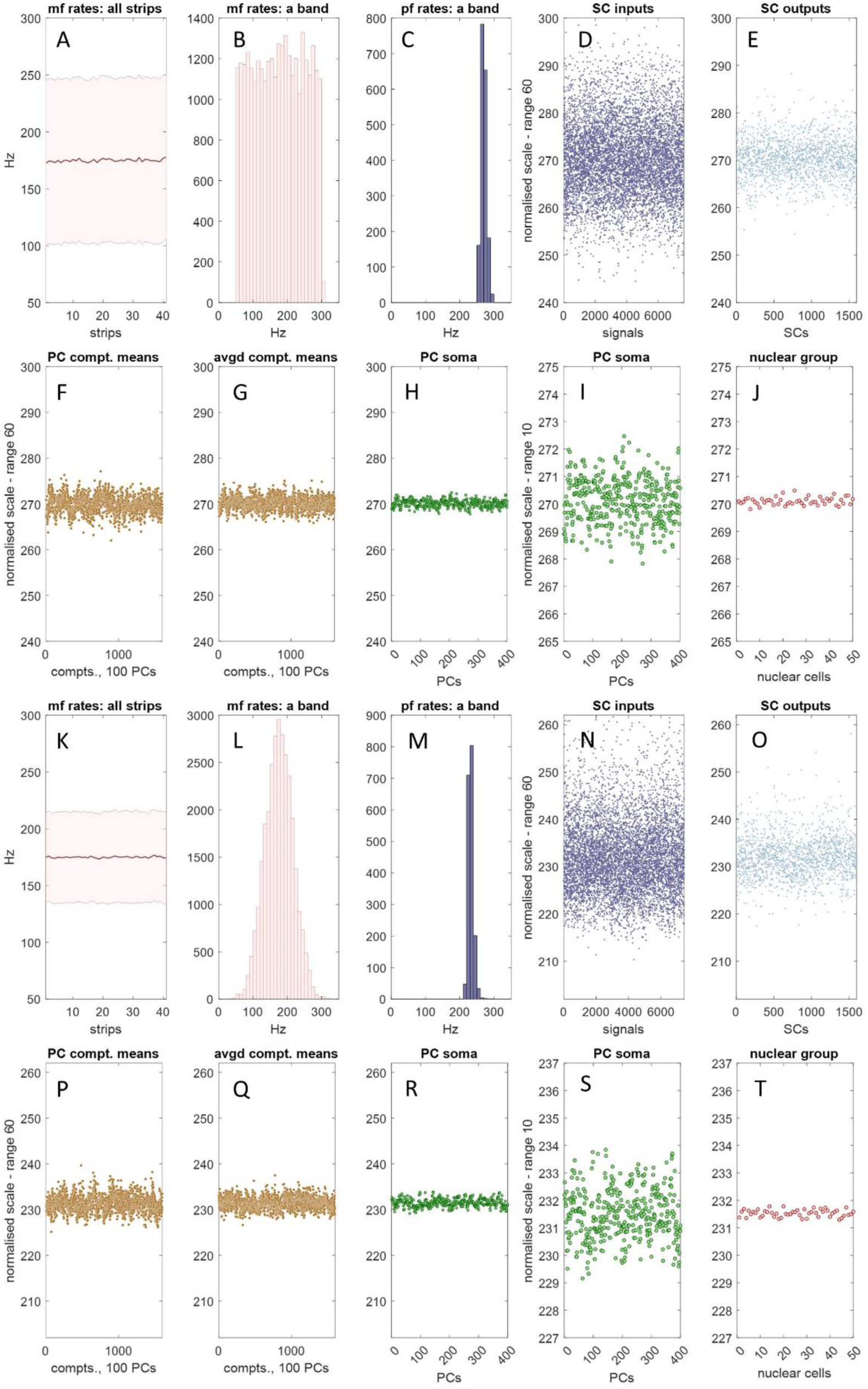
The microzone computation. SC: stellate cell; PC: Purkinje cell; compt: PC dendritic compartment; mf: mossy fibre; pf: parallel fibre; ‘rank’ and ‘file’ are defined in Fig 2. **A** Mean and SD of mossy fibre input to 41 <u>ranks</u> of the granular layer. Ranks are 150 microns wide and together make up the region supplying parallel fibre input to the microzone that equally bisects it, indicated by a dashed grey line. The morphology of mossy fibres means that all locations of a rank receive a large random sample of input rates to the whole strip. **B** Frequency distribution of mossy fibre rates received by a single <u>file</u>, 200 microns wide. Files cross ranks at right angles. Granule cell firing rates in a file are accordingly parallel fibre rates received by the middle microzone. As files cross all ranks, all middle-microzone sectors receive the same distribution of mossy fibre rates. There are 100 files. **C** Parallel fibre rates in a randomly selected file. The frequency distribution of parallel fibres rates is the same in all files. This is always true, and does not depend on ranks receiving the same mossy fibre distribution. **D** Parallel fibre rates received as input to 16 stellate cells per sector (out of 80, for illustration), obtained by randomly sampling parallel fibre rates (16 times per file, generating sample size randomly with Table 1 probabilities). The y axis range is 60 normalised Hz, down from a mossy fibre range of 250 Hz. **E** Outputs of the stellate cell unit functions which represent the cells in D. The program derives input to, and output of, all stellate cells, generating 8,000 data points, but the data are thinned for presentation. **F** We notionally divided the Purkinje cell dendritic arbour into 16 compartments per cell. Each compartment randomly samples a subset of outputs of the stellate cell unit function which represents the network of stellate cells afferent to that cell. Data are the sample means, shown for a single Purkinje cell per middle-microzone sector (so there are 1,600 values, 16 compartments x 1 Purkinje cell x 100 sectors). **G** A Purkinje cell receives inhibition from two local stellate cell networks, one on each flank. Step F was repeated, and the data averaged for each compartment. **H** Mean of the compartment means calculated for each Purkinje cell in the middle microzone (so 400 data points, 4 Purkinje cells per field x 100 fields), representing somatic integration of dendritic compartment data. **I** The same data but with the y axis range shortened to 10 normalised Hz. **J** The output of a microzone is channelled down onto a smaller cell group in the deep cerebellar nuclei, which includes the output cells of the network. Each output cell receives input from a random sample of 30–50 Purkinje cells [46]. There are ∼50 output cells in a nuclear group. Data are the sample means. **K–T** We repeated (A)–(J) but with normally distributed mossy fibre rates (SD 40), with the same result. We also obtained the same results (not shown) with a discontinuous mossy fibre distribution (range 50–300 Hz).

In this proposal, each stellate cell is the physical form of a notional unit, and the afferent data set is the frequency distribution of parallel fibre firing rates. Physically, and in the simulation, units are limited to sampling only signals in that file. Few parallel fibres are active at any time. But the high turnover of active cells means that sample size varies with time and, at any time, the number of active inputs to a unit has a distributed probability. The simulation generates sample size for each cell with the Table 1 probabilities. Firing rates received by a stellate cell are an independent random sample. The unit computation is a linear function. In physiological conditions, inputs are computed in parallel by all units (in all sectors) concurrently. All (at any moment) effectively sample the distribution of parallel fibre rates received by the whole microzone (because all sectors receive the same distribution), notwithstanding that they all receive contact from different parallel fibres. The outputs – each an analogue signal – become the data set sampled by the next layer of units.

The fast turnover of active parallel fibres means that units receive input at time-varying rates notwithstanding that granule cells typically fire in short bursts [15, 44, 45] at a rate which (as far as we know) does not vary in a burst. Accordingly (and because stellate cells fire intrinsically), unit outputs are analogue.

Stellate cell data generated by the simulation and shown in Figure 4 are the population of parallel fibre rates received as input to 16 stellate cells per sector (the number that sit between two Purkinje cells), so around 7,200 values (Fig 4D and N), and the outputs of the unit function used by the simulation to represent those cells (1,600 values, Fig 4E and O). It forms part of the proposal that a stellate cell is the physical form of the unit function.

There is direct evidence of a linear relationship of granule cell input and stellate cell rates [15] but as far as we know not confirmation, or data about the sensitivity or latency of the response. However, fidelity of transmission may be assisted. GABA spillover from stellate cell synapses enhances parallel fibre synaptic transmission to stellate cells [47, 48]. ‘It is well established that parallel fibers express functional GABA_A_Rs, that their activation depolarizes the axon, and that they increase release probability at parallel fiber synapses’ [49 p.16924, 50]. High release probability hones a rate-proportional response.

### 4.2 Purkinje cells

#### 4.2.1 PURKINJE CELL DENDRITIC COMPARTMENTS RECEIVE INNERVATION FROM AN INDEPENDENT RANDOM SAMPLE OF STELLATE CELLS AFFERENT TO THE CELL

Around 350,000 parallel fibres pass through the dendritic field of a single Purkinje cell, of which one in two make contact, normally at a single synapse (sometimes two, average 1.24) [5]. Purkinje cells are severely flattened in the plane orthogonal to parallel fibres and to the cerebellar surface, filling the molecular layer vertically, so that parallel fibres pass through by the shortest distance.

A microzone is ∼100 Purkinje cells long and 4 wide (400 is consistent with reported convergence and divergence ratios onto 50 excitatory nuclear projection cells [rats: 46]. Purkinje cells are inhibitory and fire spontaneously [40, 51–53] at a rate adjusted by simultaneous direct excitation from parallel fibres and inhibition by stellate cells.

In the simulation, a field contains 4 Purkinje cells. A Purkinje cell receives contact from stellate cells from both of its flat sides, ∼16 each side, and a smaller number of stellate cells in the two sagittally-neighbouring fields. Lateral inhibition from neighbouring fields is from a smaller number of stellate cells because not all of them extend their main axon far enough to reach, and some of them extend their main axon the wrong way. We represent those conditions as variables in the simulation. The vertical range of axon collaterals and the horizontal range of the main axon both increase with the depth of the stellate cell body in the molecular layer [41, 54]. Because of their more limited horizontal range, outer-level stellate cells are excluded in the simulation from contacting Purkinje cells in neighbouring fields.

The Purkinje cell arbour is large and branches profusely, and stellate cell contact is dispersed across it. The local postsynaptic effect is therefore a response to local inputs. We divide the Purkinje cell arbour nominally into sixteen 50 x 75 µm compartments. For morphological reasons, input to a compartment is not confined to (or even necessarily always received from) the nearest stellate cells, but each compartment is in range of most stellate cells. As a result, each compartment receives contact from a near approximation of a random subset of stellate cells afferent to a Purkinje cell.

It is unknown how many stellate cells innervate a Purkinje cell compartment. We take it as 6, around 20% of the afferent network, and that local dendritic polarisation is a linear function of the mean firing rate of stellate cells afferent to a compartment. Sampling by a compartment is independent: innervation of a compartment by a stellate cell does not exclude innervation by the same cell of other compartments. Physically, contact is limited by stellate cell shape and range to a small group of local cells. However, the stellate cell network afferent to a Purkinje cell could in theory be made up of any random sample of stellate cells in the same microzone, with the same result at population level.

#### 4.2.2 SIMULATED SAMPLING AND INTEGRATION

A notional compartment is a computational unit. Each compartment randomly samples a subset of local stellate cells. For this purpose, stellate cells are represented by outputs of the stellate cell unit function. A subset is made up of 16 of the 80 units that represent stellate cells in that sector, plus up to a further 8 in each of the sagittally neighbouring sectors. Compartment membrane voltage is a linear function of stellate cell rates afferent to that compartment. In its physical form, a compartment continually recalculates inputs (the simulation represents a moment of time). Dendritic signals are a sustained state, we propose, mitigating a randomly variable effect of spatiotemporal integration of discrete signals – a noise reduction strategy.

To declutter the data, we show unit outputs (representing Purkinje cell dendritic compartments) for a single Purkinje cell per sector (so 1,600 values in total) (Fig 4F and P). Data are by this step in a range of around 10 Hz on a normalised scale. This is down from an original mossy fibre range of 250 Hz.

A Purkinje cell receives inhibition from two stellate cell networks, one each side. We repeated the steps above for the other flanking network, then took the mean for each compartment (Fig 4G and Q).

At the next step, dendritic signals are integrated at the Purkinje cell soma. Integration is a linear function of compartment data, calculated for each Purkinje cell, in all sectors, so that this step returned 400 values (4 Purkinje cells x 100 sectors; Fig 4H, I, R and S). Modulation of somatic charge is fast and bidirectional, and reflects the average of dendritic states, which transforms into a linear effect on firing.

The result is a progressive contraction of the data range received as inputs to the system. At each step, data are coded in a frequency distribution. There is a linear relationship of the mean at each step with all other steps. This result is independent of the shape of the mossy fibre distribution (we also tested a discontinuous distribution with the same results).

Linear functions are atypical for dendritic models. What is the evidence? Purkinje cell firing is a linear function of synaptic excitation, with inhibition blocked [13, 14]. The granule cell → interneuron → Purkinje cell pathway drives a fall in the firing rate of Purkinje cells as a linear function of the strength of input from granule cells [16]. In mice during locomotion, there is a linear relationship between granule cell rates and interneuron firing, and also between excitation and inhibition of Purkinje cells and depolarising or hyperpolarising dendritic membrane voltage. Stellate cells contact Purkinje cells on smooth dendrites [41]. Smooth dendrites extend into all compartments, mitigating the effect of filtering and distance on dendritic signalling. Recording from smooth dendrites, locomotion-dependent modulation of dendritic membrane voltage ‘linearly transforms into bidirectional modulation of PC SSp [Purkinje cell simple spike] output’ [15 p.9].

The effect of the form of the parallel fibre code is to simulate simultaneous independent random sampling by single cells of the whole population of parallel fibre rates which converge on a microzone (summarised in Fig 5). This mirrors functionally simulated random sampling by granular layer units of the whole population of mossy fibre rates received as input to a rank.

**Fig 5.**
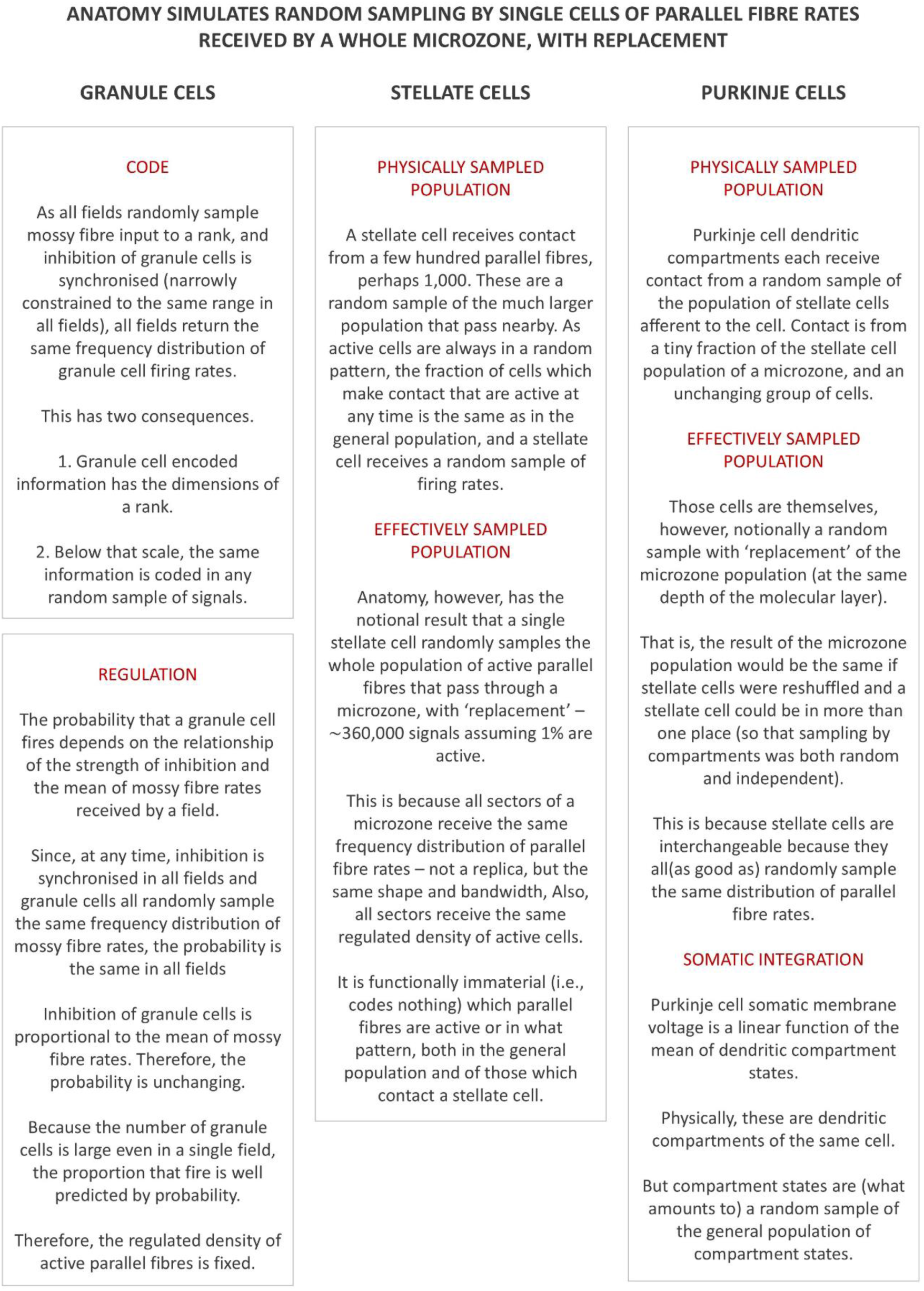
**The effect of the nature of the granule cell code on sampling in the middle layer of the cerebellar network.**

## 5. A proto-vertebrate wiring plan?

### 5.1 Recap

We have argued that the microzone computation is a computational effect of anatomy. This is not incidental but a candidate to account for cell morphologies, the three-dimensional space they occupy and the scaffold-like geometry of the connections they make. There is evidence that communication of cerebellar cells is linear [9–12]. Cells are highly adapted for faithful transmission [17–23]. Neurobiology is customised to control for biophysical and other noise. ‘Although cerebellar neurons and synapses have nonlinear intrinsic properties [55–59], these properties are disengaged or compensated in ways that keep the cerebellar network operating in a linear regime [23, 60–63]’ [24]. Firing rates linearly code metrics of behaviour [24]. Anatomy and linearity combine functionally. The result is a passive, unlearned computational effect of simply passing signals through the network.

To this point, we have a mechanism without a function. How, if at all, does a microzone discriminate between input from different ranks? Why do neighbouring microzones receive most of the same parallel fibre signals, and how is their output functionally related and coordinated?

### 5.2 Proprioception-motor loops?

The cerebellum is necessary for skilled execution and coordination of movement. We reason that in motor control, the cerebellum may process feedback to directly drive motor output.

It is probably not contentious that early vertebrate swimming, before efficient teleosts, was by undulating body movements (anguilliform). Almost the entire musculature of fish is made up of a bilaterally symmetrical row of paired axial muscle segments, termed myomeres. Movement through the water is produced by a repeating sequence of muscle contractions which cause lateral body flexion. Alternating left and right head turns set up a herring-bone pattern of high pressure which the animal rides by making itself an undulating shape.

Muscles generate analogue signals, in muscle spindles, specialised muscle fibres. Group II afferent neurons signal muscle length. Group Ia afferent neurons fire at a rate that varies with muscle length but in addition varies proportionally to the rate of muscle extension and (more modestly) inversely proportionally to the rate of muscle contraction. Say that in a vertebrate ancestor, input to a row of topographically-defined strips of the cerebellar granular layer arose from a corresponding row of contralateral myomeres, with neighbouring strips receiving the output of neighbouring myomeres. Say also that these provide drive via parallel fibres to the microzone which equally bisects the group, and that this controls motor output to the ipsilateral muscle segment opposite the middle of the row providing input. A network in this wiring plan is the junction of a proprioception-motor loop with multiple input pathways and a single output pathway. Neighbouring networks connected by parallel fibres send output to neighbouring myomeres. This connectivity is mirrored in the contralateral cerebellum, which has motor output to the other half of the body.

This is a candidate proto-vertebrate wiring plan for anguilliform swimming. Fish muscle fibres are anchored to membranes that separate myomeres. Undulations of the body cause the length of muscle fibres and the rate of change of the length of muscle fibres to vary in a rhythmically and smoothly time-varying manner. Input to a rank accordingly has a sine-wave-like signature. Input to neighbouring ranks is a small step out of phase. The size of the step depends on number of muscle segments in a full ‘cycle’ of the trunk. Passive muscle extension might in this way be used to drive muscle contractions on the other side of the body. Contractions would be automatically phase-locked and wavelength sensitive.

### 5.3 Why are loops connected by parallel fibres?

Parallel fibres coordinate motor outputs and propagate motor sequences, in this system. We briefly take these ideas in turn.

#### 5.3.1 COORDINATION OF MOTOR OUTPUTS

Cerebellar rank data are only segregated in granule cell ascending axons, while signals travel towards the cerebellar surface. Beyond that, after the granule cell axon divides in two, parallel fibre signals are randomly intermingled (within modal strata [41, 44, 64]). This means a microzone is unable to discriminate between signals from different ranks on the face of the data, compounding the cerebellar sacrifice of resolution. The sacrifice is in two steps. In the first, recoding in ranks does not discriminate between mossy fibre signals. In the second, a microzone does not discriminate between ranks.

This is not a cost but functional. It means that outputs are coordinated by topography. Immediately neighbouring microzones receive almost all the same parallel fibre input, from all the same ranks except one. Input to their next nearest neighbours differs by two ranks, and so on. We have argued that firing of Purkinje cells is a linear function of parallel fibre rates and synchronised in a microzone. Therefore, microzones receive drive which is closely coordinated with their neighbours. The relationship is closest with near neighbours and weaker with further microzones, diminishing with distance. The output of a network – a synchronised firing rate or strongly focussed firing rates of perhaps 50 nuclear output cells – is thus automatically coordinated between parallel-fibre-connected networks by the relationship of inputs.

Figure 6 illustrates **i)** input and output of a network are constrained to a phase-sensitive linear relationship, and **ii)** the relationship of the number of muscle segments in a body wave and the sensitivity of the response.

**Fig 6.**
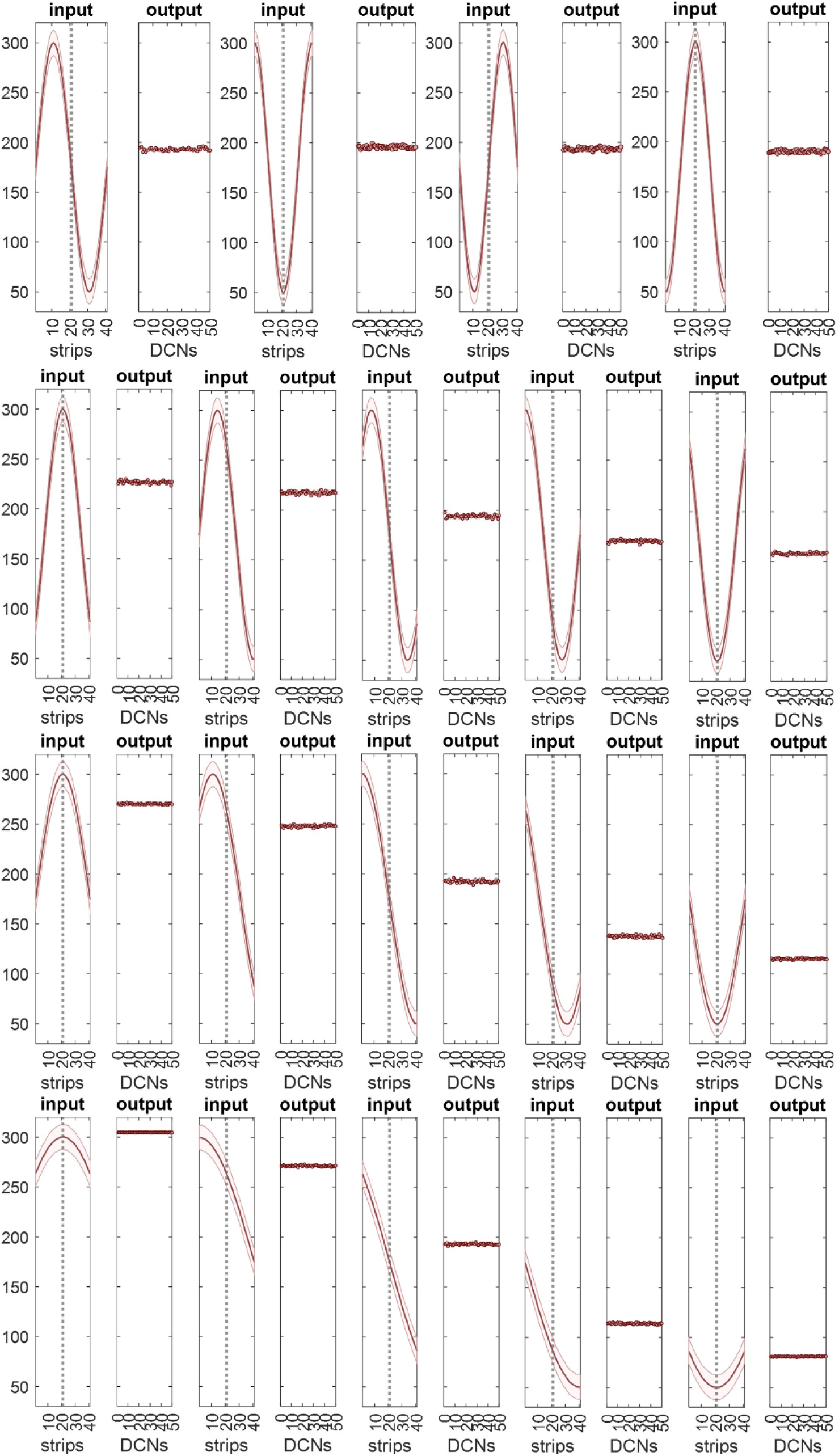
In a model of swimming, the network computation is phase sensitive. **A** Firing rates received as input to a network were varied from rank to rank, so that a plot of the mean and SD of input rates to each rank is sine-wave-like. The frequency distribution received by a single rank is normal and narrow (SD = 12.5 Hz, all ranks). The central rank is indicated by the vertical dashed line. In a locomotor cycle, a single rank receives input which varies sinusoidally with time. All ranks receive the same cycle but a step out of phase with their neighbours. Each pair of panels – 1 and 2, 3 and 4 and so on – represents a moment of time, but a different phase of the cycle, advanced 90 degrees each time. Input (to a network) is shown on the left, output on the right. Output represents firing of the output cells of the network. Output is the same in all phases: parallel fibres perfectly dampen positive feedback. **B** The same, but input rates represent 3/4 of a wavelength and the phase shift between panel pairs is 45 degrees. Output has become sensitive to cycle phase but is still dampened. **C** Input rates represent 1/2 a wavelength. The phase shift is 45 degrees. The output range is now around twice the range in (B). **D** Input rates represent 1/4 of a wavelength (45 degree phase shift). The output range increases again, now approaching the input range (and remains phase sensitive).

#### 5.3.2 SELF-DRIVING MOTOR SEQUENCES?

Motor sequences are self-driving. In anguilliform swimming, a wave of muscle contraction passes down each side of the body, 180 degrees out of phase, initiated by alternating left and right head turns. Passive extension of neck muscles contralateral to the direction of a head turn generates mossy fibre input to a corresponding row of ipsilateral ranks, and indirect input via parallel fibres to a longer row of microzones, both with strength proportional to the amount and rate of extension, with a rostro-caudal gradient. This drives motor output at related strength to a caudally-extending row of ipsilateral muscle segments, causing them to contract with proportional power. Contractions in turn cause contralateral muscle extension to propagate backwards, and so on, generating a spiral of signals generated by movement which are received as input to the cerebellum, and motor outputs.

The result of a single head turn is short-lived and ineffectual. However, alternating left and right head turns initiate backwards propagating waves of contraction – self-drive – assisted by passive undulation (modern anguilliform swimming remains in part passively powered).

## 6. Discussion

It is clear that the cerebellum is involved in learned behaviours. These can be sophisticated. When rats are trained to press a lever, the striatum receives efferent copies of cortical motor signals. Following training, the action becomes practiced and highly stereotyped. It now seems that that the basal ganglia contribute execution-related elements of the learned behaviour, such as vigour, commitment [65, 66] and kinematics [67] (vs. action selection). If cortico-striatal signals are blocked following training, performance is unimpaired [68]. If, instead, the striatum is lesioned, the learned behaviour is lost. While lesioned animals still engage with the task, they revert to their former, less practiced movements, seen early in training [67]. The same result is obtained if the pathway from the thalamus to the striatum is silenced after learning [68], indicating that the cerebellum is necessary to execute the acquired behaviour. Animals trained over a long period are able to perform the well-practiced action sequence even if the basal ganglia is then lesioned, suggesting that learned changes in the cerebellum are sufficient to execute the behaviour independently.

The changes in the cerebellum that underpin learned behaviour are not fully understood. Historically, interest has focussed on synaptic plasticity, and artificially intelligent networks often use synaptic modification to store learned changes. However, the loss and formation of synapses, growth or reduction of axons, and cell death, neurogenesis and migration, may also provide the substrate of change.

We envisage parallel fibre synaptic changes in a different role. In this view, the outcome of training under climbing fibre tuition is to polarise parallel fibre synaptic weights such that all synapses either transmit robustly (working synapses) or do not transmit at all (silent synapses). The mechanism was first proposed here [69] and is also discussed here [70]. The learning outcome is the ratio of working to silent synapses (‘synaptic ratio’). In naturally occurring conditions, silent synapses can make up a large proportion of the synaptic population [71, 72]. Working synapses are a random subset of the population. There is no pattern memory (the response does not depend on the pattern of synaptic activations). Following training, all patterns are received at the same ratio of working to silent synapses regardless of the pattern, including patterns that are not involved in training. Microzone-grouped Purkinje cells are trained and learn as a unit, so they all have the same synaptic ratio.

The function of the ratio is a gain adjustment to the anatomical computation, at network level. The adjustment is made by learned control of the number of signals that Purkinje cells receive. Because the fraction of active parallel fibres is strictly regulated, and active cells are a random (though ceaselessly changing) subset of the general population, the number is a proportion of active cells which depends on the synaptic ratio. This remains indefinitely adjustable by further training but is fixed on a behavioural timescale, and pattern-blind. All Purkinje cells in a microzone make the same adjustment. With this feature, individual gain of unit computations does not need to be taught or encoded in cell genotypes, and it can be tuned by performance.

What controls the synaptic ratio? Briefly, it depends on the relationship between parallel fibre rates received by a microzone, and the strength of excitatory drive to the inferior olive that evokes instruction signals. This is because the synaptic ratio modulates the strength of nucleo-olivary inhibitory feedback, which in turn determines whether drive evokes an instruction signal (in this model). The learning outcome is the synaptic ratio trained by the effect of feedback on the ratio of paired (with a climbing fibre signal) to unpaired parallel fibre synaptic activations.

This mechanism predicts that excitatory input to the cerebellum and to the olivary group is from the same or a related source, so that they have correctly related strength. In this contention, instruction signals are an automated part of a single system that extends outside the cerebellum (vs. executive supervision from an independent source).

## Acknowledgements

Funding: This work was supported by the Leverhulme Trust [grant number ECF-2022-079] (Mike Gilbert); the Swedish Research Council [grant number 2020-01468]; the Per-Eric och Ulla Schybergs Foundation [grant number 42630]; and the Crafoord Foundation [grant numbers 20180704, 20200729, 20220776, & 20230655] (Anders Rasmussen).

## Author contributions

M.G. researched, conceived and developed the ideas, performed the analysis, wrote the code, prepared the figures and wrote the manuscript. A.R. provided expertise and feedback, and reviewed and provided editorial comments on the manuscript.

## Notes

### Competing Interest Statement

The authors have declared no competing interest.

### Summary of Updates

The manuscript has been revised to improve the structure and the clarity of the methodology and narrative.

