## Supplementary Materials for "Computational anatomy: the cerebellar microzone computation"

SUPPLEMENTRAY MATERIALS

**1. Derivation of mossy fibre rates**

**INERVATION OF A RANK**

Each field receives a random sample mossy fibre firing rates received as input to a rank. To simulate that, we need the size of the sampled population – the number of mossy fibre signals received by a rank – and sample size, the number received by a field.

Despite the repeating ‘crystalline’ architecture of the cerebellar cortex, at local level there are anatomical random variables. Variables are: i) the number of mossy fibres that innervate a field, ii) the number of terminals they each end in, and iii) the number of those that are inside field limits.

The number of mossy fibres afferent to a field has been estimated at 100 [4]. However, that number is a field-sized fraction of the general population. The actual number is larger because mossy fibres end in a dispersed group (cluster) of terminals, and most groups straddle field limits.

We work from an estimate of the total number of terminals per field, which is, in turn, derived from convergent estimates of the number of granule cells per field. (Estimate 1) There are an estimated 1.92 x 10^6^ granule cells per μl [11]. The number of fields which fit in that volume is (1,000/150)^2^ x (1,000/200) = ~222, assuming a field depth of 150 μm [4]. The number of granule cells in a field is, therefore 1,920,000/222 = 8,649. (Estimate 2) An estimated 350,000 parallel fibres pass through a single Purkinje cell’s territory [11]. Parallel fibres run about 3 mm in each direction from the granule cell soma [12, 13]. The Purkinje cell dendritic arbour fills the molecular layer vertically and averages ~200 µm wide. Therefore, 6,000/150 = 40 fields fit in a row that provides parallel fibres that pass through a Purkinje cell dendritic territory, giving 350,000/40 = 8,750 granule cells per field. Estimates vary of the number of granule cells which extend a dendrite into a single glomerulus. 50 is mid-range. (8,750/50) x 4 = 700 glomeruli per field.

Granule cells have 3–5 dendrites, averaging 4, and receive input to each of them from a single mossy fibre (so an average granule cell receives 4 inputs). Estimates of divergence of a single mossy fibre terminal onto granule cells vary [14]. It is unclear how much this reflects biological diversity versus differences in opinion and methods. Here we use a mid-range estimate of 1:50 [15, 16]. Assuming a mossy fibre terminal receives a single granule cell dendrite from each of *d* granule cells, and given an average of 4 dendrites per granule cell and *z* granule cells per field, the total number of mossy fibre terminals in a field is *N* = (*z* x 4)/*d* = 700 given $z$ = 8,750 and $d$ = 50. A convergent estimate is provided by multiplying the per field number of mossy fibres by the average number of terminals per cluster.

Cluster size varies. We take the range as 4–12 terminals per cluster [4]. We assume the number varies at random and cluster sizes occur with equal probability. A maximum of 7 fits in a field (because a field shares the dimensions of the area that encloses a mean cluster). Fields are not anatomically bounded, so many clusters straddle field limits. To reflect that, we assume there is a 1/7 probability that a 7-terminal cluster (or larger) has 7 terminals inside a field and also the same probability of any other number. A 6-terminal cluster has a 2/7 chance that all 6 terminals are inside and a 1/7 chance of any other number, and so on.

Table 1 shows the derivation of the expected (i.e., predicted by probability) ratios of terminals contributed to a field by each cluster size (R1), not counting terminals that lie outside when a cluster straddles field boundaries; and the ratios of mossy fibres contributing *n* = 1–7 terminals (R2), regardless of cluster size. R1 ratios are given by the sum of numbers in each row. So, for example, the ratio of cells that end in 8 terminals to cells that end in 5 is 28:25. R2 ratios are given by the number of appearances of each number in the grid – so the expected ratio of cells which contribute 7 terminals to the number which contribute 4 is 6:12. The ratios effectively give the distributed probability of possible states. For example, there is a probability of 10/(6 + 8 + 10 + 12 + (3 x 9)) = 0.159 that a mossy fibre contributes 5 terminals to a field (regardless of cluster size). Similarly, there is a probability of ((6 x 28)/((6 x 28) + 27 + 25 + 22))/7 = 0.099 that a given cell contributes 7 terminals to a field.

**Table 1. Derivation of expected ratios**

| **size** | **number of terminals inside** | | | | | | | **R1** |
| --- | --- | --- | --- | --- | --- | --- | --- | --- |
| **12** | 7 | 6 | 5 | 4 | 3 | 2 | 1 | **28** |
| **11** | 7 | 6 | 5 | 4 | 3 | 2 | 1 | **28** |
| **10** | 7 | 6 | 5 | 4 | 3 | 2 | 1 | **28** |
| **9** | 7 | 6 | 5 | 4 | 3 | 2 | 1 | **28** |
| **8** | 7 | 6 | 5 | 4 | 3 | 2 | 1 | **28** |
| **7** | 7 | 6 | 5 | 4 | 3 | 2 | 1 | **28** |
| **6** | 6 | 6 | 5 | 4 | 3 | 2 | 1 | **27** |
| **5** | 5 | 5 | 5 | 4 | 3 | 2 | 1 | **25** |
| **4** | 4 | 4 | 4 | 4 | 3 | 2 | 1 | **22** |
| **R2** | 6 | 8 | 10 | 12 | 9 | 9 | 9 |  |

If R1 ratios are denoted by a series $x_{4},x_{5}\ldots x_{12},$ and a given element in the series is $x_{j}$, the fraction of terminals provided by a given cluster size is $x_{j}/\left( x_{4}+x_{5}+\ldots+x_{12} \right)$, and this fraction is multiplied by $N$ to give the number of terminals. The number of terminals is divided by the expected mean number of terminals that fall inside field limits (for that cluster size) to give the expected number of mossy fibres that provide them. The number of $j$-terminal mossy fibres that innervate a field is therefore given by

$$N\left[ \frac{x_{j}}{\sum_{i=1}^{n} x_{i}} \right]/\left[ \frac{x_{j}}{7} \right]$$

The results are shown in Table 2. The equal number of mossy fibres (mfs) in each column is what we would expect, because we made it an assumption that all sizes occur with equal probability, and mf values are expected values – i.e., in proportions predicted by their probability. The mf total is the expected number of mossy fibres afferent to a field (which contribute at least one terminal).

**Table 2. The expected number of mossy fibres which innervate a field**

| **cluster size** | **12** | **11** | **10** | **9** | **8** | **7** | **6** | **5** | **4** | **total** |
| --- | --- | --- | --- | --- | --- | --- | --- | --- | --- | --- |
| **terminals** | 81 | 81 | 81 | 81 | 81 | 81 | 78.1 | 72.3 | 63.6 | 700 |
| **mfs** | 20.25 | 20.25 | 20.25 | 20.25 | 20.25 | 20.25 | 20.25 | 20.25 | 20.25 | 182.25 |

R2 ratios – the relative incidence of afferent mossy fibres that contribute 7 terminals to a field, 6 terminals, 5 and so on, down to 1 – and the expected number of mossy fibres that contribute them, are shown in Table 3.

**Table 3. The number of mossy fibres which contribute *n* = 1–7 terminals to a field**

| **n** | **7** | **6** | **5** | **4** | **3** | **2** | **1** | **total** |
| --- | --- | --- | --- | --- | --- | --- | --- | --- |
| **R2** | 6 | 8 | 10 | 12 | 9 | 9 | 9 | 63 |
| **R3** | 42 | 48 | 50 | 48 | 27 | 18 | 9 |  |
| **mfs** | 17.34 | 23.15 | 28.92 | 34.72 | 26.04 | 26.04 | 26.04 | 182.25 |
| **%age** | 9.52 | 12.69 | 15.87 | 19.05 | 14.29 | 14.29 | 14.29 |  |

R3 is *n* x R2, giving the ratio of terminals provided by mossy fibres with *n* terminals inside a field. %age is given by (R2/63) x 100. this is the percentage of mossy fibres with *n* terminals inside a field.

From here, the number of mossy fibres that innervate a rank is given by 182.25 x 100 divided by the mean number of clusters per mossy fibre per rank. If a rank receives, on average, five clusters per mossy fibre, ~3,645 mossy fibres innervate a rank. This is the estimate used in the simulation unless stated otherwise.

*Assumptions.* Cluster sizes occur with equal probability. The probabilities of each outcome when a cluster straddles field boundaries (how many terminals are inside and how many outside) are as stated. A single mossy fibre ends in an average of 5 terminal clusters per rank.

**INNERVATION OF A FIELD**

Terminal clustering means a field receives multiple copies of most signals. The proportions of mossy fibres afferent to a simulated field that contribute each possible number of terminals (1–7) are generated in simulated steps. In the first step, the simulation randomly generates, for each field, the cluster size ratio. This means, the relative proportions of afferent mossy fibres (which contribute at least 1 terminal to a field) that end in 4 terminals, 5, 6 and so on up to 12, assuming they occur with equal probability, disregarding how many terminals fall inside and outside field boundaries. In step two, the ratio of cluster sizes is weighted to reflect the greater probability that large clusters contribute more terminals to a field. In step 3, ratios are converted to the fraction of terminals contributed by each cluster size. When multiplied by the total number of terminals, this gives the number of terminals contributed by each cluster size. In step four, the values obtained in step three are divided up between cells that contribute a single terminal, 2 terminals, 3 and so on, to reflect straddling of field limits. The proportions give the number of mossy fibres afferent to a simulated field that contribute each number of terminals up to seven, the maximum.

There is no field-level effect of mossy fibre terminal branching. Terminal clusters arising from the same mossy fibre are separated by a minimum physiological distance, so that a field cannot contain terminals of more than one cluster per mossy fibre. Fields receive a randomly variable number of copies of most signals in the cerebellum and the simulation. In the simulation, copies are added after sampling of mossy fibre rates to obtain rates received by a field. An alternative order would be to add copies to the sampled population and then sample that. We follow anatomy: the number of *rates* received by a field is equal to the number of afferent mossy fibres; the number of *signals* contains duplicate rates.

**2. Evidence of linear transmission in the granular layer**

**LINEAR TRANSMISSION OF MOSSY FIBRES TO GOLGI CELLS**

Golgi cells are large interneurons whose cell bodies and basal dendrites lie in the granular layer. Mossy fibres directly contact Golgi cell basal dendrites. Golgi cell firing frequency increases linearly with the amplitude of depolarising current [17 p.845]. ‘Golgi cells can follow peripheral signals in a continuous fashion, modulating their frequency with the intensity of the stimulus [citing [18, 19]]’ [20]. As a result, during movement, the ‘output of Golgi cells varies smoothly as a function of input.…They are capable of modulating their output to reflect their input with little noise or signal distortion’ [21]. ‘Sensory-evoked Golgi-cell inhibition scales proportionally with the level of mossy fiber excitatory synaptic input’, such that inhibition reliably conveys mossy fibre rate information [22]. On this evidence: mossy fibre rate information is conserved proportionally in Golgi cell firing rates.

Golgi cells extend their basal dendrites into glomeruli [23, 24]. Contact on them by mossy fibres is multi-synaptic [25], contributing to a reliable, rapid (submillisecond) response. The fast rise time and weak distance dependence of the amplitude and timing of Golgi cell EPSCs evoked by mossy fibre signals, suggests that basal dendrites introduce relatively little filtering [26, 27], as expected for large-diameter dendrites. We note that the pause in Golgi cell firing following a discrete stimulus under anaesthesia [22, 28, 29] disappears from Golgi cell recordings during locomotion [19].

During behaviour in freely-moving animals, mossy fibre activity is dense: a high fraction of mossy fibres are active [30]. Both mossy fibres [31-33] and Golgi cells [19] have been reported to fire with a sustained, time-varying signature. In the behaving animal, Golgi cell basal dendritic membrane potential is a continuous variable under modulation by sustained inputs, we submit. As charge transfer to the soma is passive, polarisation of the soma is likewise sustained, under modulation by dendritic states.

It is worth noting that very high mossy fibre rates can be generated in experimental conditions. However, the typical physiological range is 50–300 Hz [32]. In the simulations, we use the physiological rates, within stated constraints on the shape of the distribution.

**IS THERE OTHER CONTROL OF GOLGI CELLS?**

Do Golgi cells receive a significant influence of input from other sources? The evidence is incomplete but currently suggests other influence is absent or weak. Contrary to early reports, neither Purkinje cells nor climbing fibres contact Golgi cells [34, who give references], and only a modest minority of inhibitory inputs to Golgi cells (if any) are from molecular layer interneurons [35], which generate weak synaptic currents [36], consistent with either extremely weak or wholly absent innervation [37]. There is conflicting evidence whether Golgi cells make inhibitory synaptic contact on each other [37, 38]. Our simulation does not include an effect on Golgi cell firing by other sources of input.

**GOLGI CELL OSCILLATIONS**

Golgi cells fire autonomously. Under anaesthesia, firing falls into a slow oscillating pattern [39]. Under excitatory synaptic input, however, this disappears [38]. Anaesthesia ‘has a strong influence on spontaneous activity of Golgi cells’ [34]. Discharge at low rates with irregular timing under anaesthesia [40] is replaced by higher rates with more regular timing without anaesthesia [18, 19, 41]. This paragraph is a cautionary note about evidence obtained under anaesthesia, which has been used to argue that oscillations provide a form of signalling.

**LINEAR TRANSMISSION OF MOSSY FIBRES TO GRANULE CELLS**

There is significant evidence that granule cell firing rates are linearly related to input rates, but also evidence that has been given a conflicting interpretation. We take them in turn.

The short and equal length of granule cell dendrites would suggest light and equal filtering. The mossy fibre-granule cell connection has a range of adaptations which support high-fidelity transmission of high frequency signals across a wide bandwidth [42]. Faithful transmission of rate information is reported [14, 43]. Vesicle release and replenishment are fast [44, 45] and postsynaptic AMPA receptors operate in their linear range [45], where they are resistant to desensitisation [46]. Multiple contacts are made by a mossy fibre with each granule cell [16]. Conversion of depolarising somatic charge into granule cell firing is a simple function, such that granule cells ‘have a relatively linear and uncomplicated conversion of depolarisation level to spike rate’ [47 p.2393, citing Jorntell and Ekerot 2006 and D'Angelo et al 1998].

Glutamate spillover enhances precision and reliability of transmission [45, 48]. During behaviour, at sustained physiological mossy fibre rates, there may be a build-up of intraglomerular glutamate, increasing the relative influence of spillover in a balance with synaptic transmission, assisted by short-term synaptic depression. In these conditions, spillover may dominate, as it dominates inhibitory transmission [49, 50]. Spillover from multiple release sites (200–400 at a single mossy fibre terminal) is plausibly sufficient to mitigate variability of vesicular size, release probability, and the number of release points at synapses made on any individual cell, increasing fidelity and equality of excitation of granule cells.

Against that, there are reported to be heterogeneous mossy fibre-granule cell synaptic weights. Different strength and short-term dynamics of the mossy fibre to granule cell connection have been reported in vestibular slices, with GABA_A_ receptors blocked. The amplitude of the response, measured at the soma, is reported to be statistically correlated to the source of the mossy fibre that is stimulated [51], suggesting that the strength of the connection depends on the source.

However, the presence of weights does not necessarily mean that the function is to modulate firing rates. Rather, the authors suggest that the function may temporally code the source of inputs because different combinations have different response onset times. An alternative, which they also mention, is that some inputs may be in a supporting role, reminiscent of ‘driver’ and ‘modulatory’ signals in the thalamus and cortex [52]. Feasibly, signals in a supporting role must be present for the postsynaptic granule cell to fire, but do not affect the granule cell firing rate.

**LINEAR TRANSMISSION OF GOLGI CELLS TO GRANULE CELLS**

The granular layer hypothesis includes the proposal that inhibition of granule cells is, at any time, linearly related to the mean firing rate of Golgi cells afferent to each glomerulus. In support, we cite evidence that GABA spillover into the intraglomerular space is proportional to afferent Golgi cell rates and that inhibition of granule cells is almost exclusively mediated by spillover, and therefore linearly reflects the intraglomerular concentration of GABA.

Fine, beaded axon fibres enter glomeruli, where they inhibit granule cells [24]. ‘In the adult rat cerebellum, each granule cell dendrite receives 2.6 ± 0.55 synaptic contacts from Golgi axon terminals’ [53] citing [16].^[1]^ However, the large majority of inhibition (98%) is by spillover [49], where neurotransmitter released into the synaptic cleft spills out into the glomerular space. Golgi cells release GABA, an inhibitory neurotransmitter. This is detected by high-affinity GABA_A_ receptors located perisynaptically and extrasynaptically on granule cells [54, 55]. Even synaptically-received signals have most of their effect (i.e., the large majority of charge transfer is mediated) via spillover.^[2]^

Golgi cells fire [40] at a time-varying rate in the behaving animal, so that a glomerulus receives continuous input. As a result, there is a sustained build-up of glomerular GABA during behaviour [56] at an adjustable concentration controlled by Golgi cell firing rates [50]. Signalling by spillover is sometimes assumed to be ambient and slow. However, the action of glomerular GABA spillover has a fast phasic component – not as fast as synaptic transmission (~1 ms) but with a rise time of only a few milliseconds [50]. Unlike the spiky appearance of synaptically-induced inhibitory postsynaptic currents, spillover [22] generates a sustained outward current.
